# Ceramide synthase 6 (CerS6) promotes alcohol-induced fatty liver by promoting metabolic dysfunction and upregulating lipid droplet-associated proteins

**DOI:** 10.1101/2022.11.03.514971

**Authors:** Sookyoung Jeon, Eleonora Scorletti, Delfin Buyco, Chelsea Lin, Yedidya Saiman, Jasmin Martin, Royce Hooks, Besim Ogretmen, Josepmaria Argemi, Luma Melo, Ramon Bataller, Rotonya M. Carr

**Author notes:** **Corresponding author:** Rotonya M. Carr, MD, FACP, Address: Department of Medicine, Division of Gastroenterology, Box 356424, 1959 NE Pacific Street, Seattle, WA 98195-6424.

## Abstract

**Objective:** Alcohol-associated liver disease (ALD) is the leading cause of liver-related mortality worldwide. Current strategies to manage ALD largely focus on advanced stage disease, however, metabolic changes such as glucose intolerance are apparent at the earliest stage of alcoholic steatosis and increase the risk of disease progression. Ceramides impair insulin signaling and accumulate in ALD, and metabolic pathways involving ceramide synthase 6 (CerS6) are perturbed in ALD during hepatic steatosis. In this study, we aimed to investigate the role of CerS6 in ALD development.

**Methods:** C57BL/6 WT and CerS6 KO mice of both sexes were fed either a Lieber-DeCarli control (CON) or 15% ethanol (EtOH) diet for 6 weeks. *In vivo* metabolic tests including glucose and insulin tolerance tests (GTT and ITT) were performed. The mice were euthanized, and liver histology and lipid levels in serum and liver were measured. For *in vitro* studies, CerS6 was deleted in human hepatocytes and were incubated with EtOH and/or C_16:0_-ceramides. RNAseq analysis was performed in mice and in liver from patients with different stages of ALD and diseased controls.

**Results:** After six weeks on an EtOH diet, CerS6 KO mice had reduced body weight, food intake, and %fat mass compared to WT mice. Male (but not female) EtOH-fed KO mice showed significantly higher O_2_ consumption, CO_2_ production, respiratory exchange ratio, and energy expenditure (*P*<0.05 for all) during the dark period compared to EtOH-fed WT mice. In response to EtOH, WT mice developed mild hepatic steatosis, while steatosis was alleviated in KO mice as determined by H&E and ORO staining. KO mice showed significantly decreased long-chain ceramide species, especially C16:0 ceramides, in the serum and liver tissues compared to WT mice. CerS6 deletion decreased serum TG and NEFA only in male not female mice. CerS6 deletion improved glucose tolerance and insulin resistance in EtOH-fed mice of both sexes. RNAseq analysis revealed that 74 genes are significantly upregulated and 66 genes are downregulated by CerS6 deletion in EtOH-fed male mice, with key network pathways including TG biosynthetic process, positive regulation of lipid localization, and fat cell differentiation. Similar to RNAseq results, absence of CerS6 significantly decreased mRNA expression of lipid droplet associated proteins in EtOH-fed mice. *In vitro*, EtOH stimulation significantly increased PLIN2 protein expression in VL-17A cells while CerS6 deletion inhibited EtOH-mediated PLIN2 upregulation. C_16:0_-ceramide treatment significantly increased PLIN2 protein expression compared to CON. Importantly, progression of ALD in humans was associated with increased CerS6 hepatic expression.

**Conclusions:** Our findings demonstrate that CerS6 deletion attenuates EtOH-induced weight gain and hepatic steatosis and improves glucose homeostasis in mice fed an EtOH diet. Notably, we unveil that CerS6 plays a major role as a regulator of lipid droplet biogenesis in alcoholic intra-hepatic lipid droplet formation. Together, our data suggest that CerS6 may be targeted for treatment for early stage ALD.

## 1. Introduction

Alcohol-associated liver disease (ALD) is a leading cause of liver-related morbidity and mortality worldwide [1]. Despite a need for effective therapeutics for ALD, there are no approved drugs. In ALD, hepatic lipid accumulation is the earliest and most common response to chronic alcohol consumption. Simple steatosis is asymptomatic and can be reversible with alcohol cessation. But prolonged steatosis can further progress to advanced stages of liver injury such as steatohepatitis, hepatic fibrosis, and cirrhosis, and this progression is associated with insulin resistance [2, 3]. Insulin resistance favors the progression of ALD to cirrhosis [4]. Previously, we have demonstrated that perilipin 2 (PLIN2), a major hepatocellular lipid droplet (LD) protein on the LD surface, is required for the development of steatosis and insulin resistance in ALD [5], and that PLIN2 is positively associated with ceramide synthesis [6].

Ceramide, as a precursor for all sphingolipids, resides in the center of sphingolipid metabolism. Structurally, ceramide consists of a sphingosine base and an amide-linked fatty acyl chain of varied length within typically 14 to 26 carbons. Ceramides are generated by three pathways: 1) the *de novo* synthetic pathway, 2) the sphingomyelin hydrolysis pathway, and 3) the salvage/recycling pathway. Ceramide synthases (CerSs) are a group of ceramide synthesizing enzymes involved in both *de novo* and salvage pathways. Among six mammalian CerSs (CerS1–CerS6), CerS6 has a substrate preference for C_16:0_-ceramide [7]. CerS6 is ubiquitously expressed across various tissues at basal levels, [8] and can be upregulated, producing C_16:0_-ceramides in response to metabolic insults such as alcohol or obesogenic diet [9, 10].

Among alcohol-induced dysregulated lipid pathways [11], aberrant ceramide metabolism has emerged as an important pathological feature of ALD. Ceramides are bioactive sphingolipids that can impair insulin action and induce inflammation and mitochondrial dysfunction [12]. Hepatic ceramide levels are increased in ALD patients and murine models [10, 13]. Moreover, enzymes involved in ceramide synthesis such as CerSs and serine palmitoyl transferase (SPT) are dysregulated in ALD patients [13]. In addition, pharmacological and genetic inhibition of overall ceramide generation have shown to ameliorate alcohol-induced steatosis in alcohol-fed animals [10, 14-16]. Previously, we and others demonstrated that CerS6 has pathological functions in several metabolic diseases including obesity, NAFLD, and ALD [9, 10]. Furthermore, Sociale et al. suggested that Drosophila CerS isoform has dual functions as a ceramide-producing enzyme as well as a transcription regulator [17], both of which are potentially implicated in ALD pathogenesis. Despite these findings, little is known about the role of CerS6 and its main product C_16:0_-ceramides in the onset of ALD development. In this study, we explored if genetic inhibition of CerS6, the enzyme involved in ceramide *de novo* synthesis and salvage pathways, can alleviate EtOH-induced hepatic steatosis and insulin resistance.

## 2. Material and Methods

### 2.1. Animal models

All animal procedures were conducted in compliance with protocols approved by the Institutional Animal Care and Use Committee of the University of Pennsylvania. All efforts were made to minimize animal discomfort. CerS6-deficient mice on a C57Bl/6 background were provided by Dr. Besim Ogretmen at Medical University of South Carolina, and CerS6 knockout (KO) animals were maintained through heterozygous breeding. Wild-type (WT) C57Bl/6 mice were purchased from Jackson laboratory. Genomic tail DNA was used to genotype offspring by PCR. The primers were as follows: Reverse: 5′-CACACCCATATGGAACTCTTACA-3′; Forward1: 5′-TTCGGTTAAGAATGGCCTTG-3′; and Forward 2: 5′-CCAATAAACCCTCTTGCAGTTGC-3′. PCR reaction conditions were 94°C for 2 minutes, 10 cycles of 94°C for 15 sec, 64,2/51.2°C for 30 sec, and 72°C for 30 sec, then 30 cycles of 94°C for 15 sec, 58°C for 1 min, and 72°C for 30 seconds. Expected PCR products are 295□bp for CerS6 KO, 460□bp for wild type, or both for heterozygous mice (**Supplementary Figure 1A**). Protein knockout was confirmed in the liver and adipose tissue via western blotting (**Supplementary Figure 1B**).

For animal studies,10-11 wk old WT and CerS6 KO male and female mice received either Lieber-DeCarli control (CON) or 15% ethanol (EtOH) diet (Bioserv, Flemington, NJ) for 6 weeks. The CON Lieber-DeCarli diet has 35.9% fat, 49% carbohydrates, and 15.1% protein calorie content. The ethanol-containing diet has ethanol added to account for 15% of total calories with the equivalent caloric amount of carbohydrates removed. To acclimate mice to the EtOH diet, mice were given 5% ethanol calorie content for two days, then 10% ethanol calorie content for two days prior to start of the 15% ethanol calorie content diet. Mice were allowed ad libitum access to food and water and maintained in a facility with a 12 hr light/dark cycle at 22 ± 2°C. Food intake and body weight were measured daily and three times per week, respectively. After dietary treatment, mice were anesthetized with isoflurane before collecting retro-orbital blood samples. Tissues were harvested, immediately frozen in liquid nitrogen, and stored at −80°C for further analyses.

### 2.2. ITT/ GTT

Insulin tolerance test (ITT) and glucose tolerance test (GTT) were performed at 6 weeks in EtOH-fed WT and CerS6 KO mice. For the GTT, animals were fasted for five hours before 2 g/kg glucose solution was intraperitoneally administered. For the ITT, animals were fasted five hours before 0.75U/kg of human insulin was intraperitoneally administered. Tail blood glucose was measured at time 0 (before glucose injection), 15, 30, 60, and 90 minutes with a glucometer (Lifescan, Inc., Milipitas, CA). Mice were allowed to recover and resume their diets after completion of the testing.

### 2.3. Metabolic analyses

Energy expenditure (EE) was measured by indirect calorimetry using a Comprehensive Laboratory Animal Monitoring System (Columbus Instruments, Columbus, OH). Following a 24 h acclimation period, O_2_ consumption, CO_2_ production, energy expenditure, and locomotor activity (measured by x, y and z axis infrared beam breaks) were determined. Respiratory exchange ratio (RER) was calculated by dividing the volume of CO_2_ produced by the volume of O_2_ consumed (VCO_2_/VO_2_). Fat and lean mass were measured in non-fasting mice by using nuclear magnetic resonance spectroscopy (EchoMRI-100; Houston, TX) at six weeks.

### 2.4. Serum and liver biochemical assays

Serum triglycerides (TGs; Stanbio, St. Boerne, TX), alanine aminotransferase (ALT; Stanbio), and non-esterified FAs (NEFAs; Wako, Richmond, VA) were measured using enzymatic colorimetric assays according to the manufacturer’s instructions. Lipids were extracted from livers for TG measurement as previously described [16], and hepatic TGs were measured using commercial kits (Stanbio).

### 2.5. Sphingolipid and fatty acid analyses

For hepatic sphingolipid and fatty acid analyses, liver was homogenized in RIPA buffer and protein levels were quantified using the Pierce™ BCA protein assay kit (Thermo Fisher Scientific, Waltham, MA). Liver lysate and serum samples were analyzed by mass spectrometry for sphingolipid and fatty acid content at the metabolomics core at the Medical University of South Carolina. Results are normalized per milligram protein in the liver lysate.

### 2.6. Histology

Dissected liver samples were placed in a cartridge, fixed in 10% buffered formalin overnight, and then transferred to 70% EtOH until paraffin embedding. Paraffin sections were stained with hematoxylin and eosin (H&E). To visualize lipid deposition in the liver, liver sections frozen in cryoprotectant media were stained with Oil-red-O (ORO). Sectioning and staining of the liver were performed by the Molecular Pathology and Imaging Core at the University of Pennsylvania. Slides were visualized under bright field with Nikon 80i microscope and images were captured with a Nikon DS-Qi1MC camera and image analysis system (Nikon Instruments, Melville, NY).

### 2.7. In vitro studies

VL17A cells were a kind gift from Dr. Dahn Clemens at the University of Nebraska. CerS6 knockout VL17A cell line was established using the CRISPR-Cas9 system from Synthego (Synthego Corporation, Menlo Park, CA, USA). Successful deletion of CerS6 was confirmed by immunoblotting. Cells were grown under standard cell culture condition at 37°C in Dulbecco’s Modified Eagle’s Medium supplemented with 10% fetal bovine serum and 1% penicillin/streptomycin. 1 × 10^6^ CerS6 CRISPR-knockout or CRISPR-control VL17A cells were plated onto 6-well plate. Cells were incubated with standard media, or supplemented with either ethanol (100mM), C_16:0_-ceramide (100nM) (Cayman Chemical, Ann Arbor, MI), or ethanol and ceramide (100mM and 100nM, respectively) for 24hr. Cell viability was assessed by CellTiter 96® Aqueous One Solution Cell Proliferation Assay (Promega, Madison, WI) according to manufacturer’s instructions.

### 2.8. Immunoblot analyses

Liver or cell lysates were prepared in ice-cold RIPA buffer supplemented with protease inhibitor cocktail and phosphatase inhibitor cocktail tablets (Roche Diagnostics, Mannheim, Germany). Protein concentration was assessed by Pierce BCA assay (Thermo Fisher Scientific). 50 μg total protein was loaded in each well. Samples were separated by SDS-PAGE and transferred onto PVDF membrane. Blots were blocked for 1hr in 5% non-fat dry milk in Tris-buffered saline Tween 20 (TBST) followed by overnight incubation at 4°C with antibodies against Plin2 (1,000; Abcam), Plin3 (1:000, Abcam), and CerS6 (1,000, Abnova). GAPDH (1,000, Cell Signaling) was used as an internal control. After overnight incubation, blots were washed 3x in TBST for 10 min each and were then incubated with anti-rabbit secondary antibody (1:5000, Santa Cruz) or anti-mouse secondary antibody (1:10000, Santa Cruz) at room temperature for 1hr. All blots were visualized by chemiluminescence and quantified by using ImageJ software (NIH, Bethesda, MD).

### 2.9. Quantitative real time PCR analysis

RNA was extracted using the PureLink™ RNA mini kit (Life Technologies, Carlsbad, CA). RNA was treated with DNase I (Life Technologies), and reverse transcribed using a high-capacity cDNA reverse transcription kit (Applied Biosystems). mRNA expression was measured by real-time PCR using either SYBR Green primers (IDT, Coralville, IA) or TaqMan primers (Life Technologies). SYBR Green Primer sequences were obtained from Primer Bank (http://pga.mgh.harvard.edu/primerbank/citation.html). Relative mRNA expression was normalized to GAPDH, 18s rRNA, or 36B4.

### 2.10. RNA-seq

Total RNA quantity and quality were assayed with an Agilent 2100 bioanalyzer instrument using the RNA 6000 Nano kit (Agilent Technologies). Libraries were prepared at Next Generation Sequencing Core at the University of Pennsylvania using TruSeq Stranded mRNA HT Sample Prep Kit (Illumina) as per the standard protocol in the kit’s sample preparation guide. Libraries were assayed for size using a DNA 1000 kit of Agilent 2100 Bioanalyzer (Agilent Technologies) and quantified using the KAPA Library quantification kit for Illumina platforms (KAPA Biosystems). One hundred base pair single-read sequencing of multiplexed samples was performed on an Illumina HiSeq 4000 sequencer. Illumina’s bcl2fastq version 2.20.0.422 software was used to convert bcl to fastq files.

Penn Genomic Analysis Core at University of Pennsylvania performed data analysis. Raw sequence files (fastq) for 20 samples were mapped using salmon (https://combine-lab.github.io/salmon/) against the mouse transcripts described in genecode (version vM25, built on the human genome GRCm38, https://www.gencodegenes.org), with a 90.7% average mapping rate yielding 17.0M average total input reads per sample. Transcript counts were summarized to the gene level using tximport (https://bioconductor.org/packages/release/bioc/html/tximport.html), and normalized and tested for differential expression using DESeq2 (https://bioconductor.org/packages/release/bioc/html/DESeq2.html). Statistical results for pairwise contrasts of interest, overall genotype, overall diet and the interaction between diet and genotype were exported. Clustered heat-map visualizations were generated with ClustViz (https://biit.cs.ut.ee/clustvis/), based on vst-normalized expression values exported from DESeq2. Volcano plot was made on GraphPad, and gene enrichment analysis was performed using Metascape (https://metascape.org).

### 2.11. Human studies

For human RNA-seq studies, details are described in Argemi et al. [18]. Human liver samples were obtained from the Human Biorepository Core from the National Institutes of Health-funded international InTeam consortium and from Cliniques Universitaires Saint-Luc (Brussels, Belgium). All participants gave written informed consent, and the research protocols were approved by their local Ethics Committees and by the central Institutional Review Board of the University of North Carolina at Chapel Hill. A total of 52 subjects were grouped into seven categories; non-obese, high alcohol intake, early alcoholic steatohepatitis (*N* = 12); non-severe alcoholic hepatitis (*N* = 11); severe alcoholic hepatitis responders to medical therapy (*N* = 9); severe alcoholic hepatitis non-responders to medical therapy (*N* = 9), explants from patients with AH (*N* = 11).

### 2.12. Statistics

Statistical analyses were performed using GraphPad Prism 8.2 (San Diego, CA, USA). All data were presented as mean ± standard error of measurement (SEM). Statistical analysis was performed using t-test or one-way ANOVA with Tukey’s multiple comparisons test. Statistical significance was determined at p<0.05.

## 3. Results

### 3.1. Effects of CerS6 ablation on body weight, food consumption, and adiposity

To investigate the role of CerS6 in ALD pathogenesis, we performed *in vivo* studies using WT and CerS6 whole-body KO mice. Following CON and EtOH feeding for 6 weeks, various measurements were assessed in WT and CerS6 KO mice of both sexes. Here, we primarily focused on comparing WT and CerS6 KO mice fed an EtOH-diet. In EtOH-fed male and female mice, body mass gradually increased over the feeding period (**Figure 1A**). There were no significant body weight changes between genotype matched CON-fed and EtOH-fed mice (**Supplementary Figure 2**). After 14 days of EtOH feeding, CerS6 KO mice exhibited significantly less body weight gain than the WT mice with less food consumption. Similar to male mice, female CerS6 KO mice showed significantly less body weight gain over the feeding period (except week 6) with less food consumption compared to the WT mice (**Figure 1A, B**). In both male and female groups, CerS6 KO mice showed significantly lower %fat mass, and %epididymal white adipose tissue (eWAT) mass compared to the WT mice (**Figure 1C, D**). In both sexes, liver weight was significantly reduced in the EtOH-fed CerS6 KO mice compared to WT mice (**Figure 1E**).

**Figure 1.**
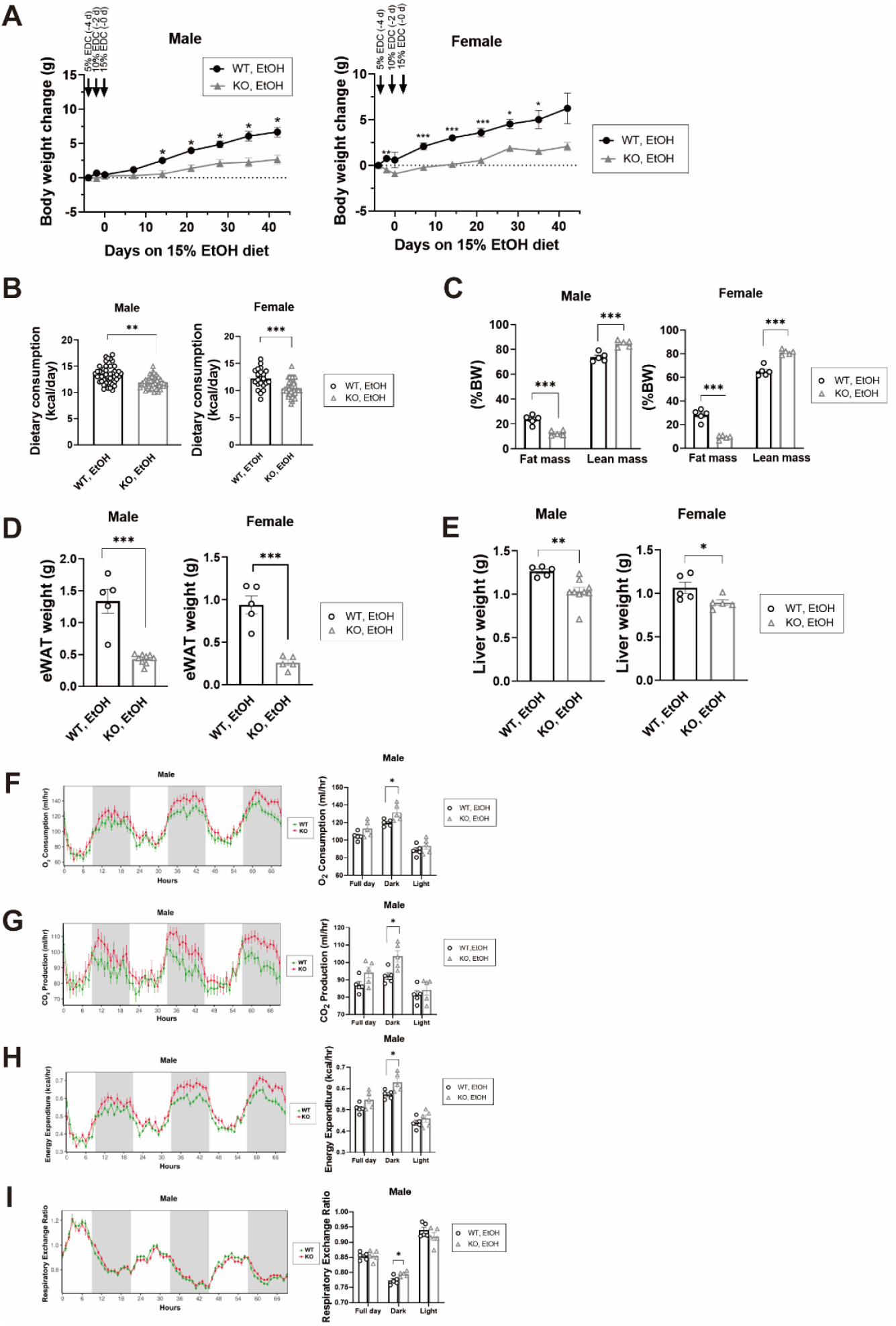
CerS6 ablation reduces body weight gain (A), cumulative dietary consumption (B), adiposity (C), epididymal white adipose tissue (eWAT) weight (D), and liver weight (E), and alters metabolic phenotyping (F-I) in the EtOH-fed mice. WT and CerS6 KO male and female mice were fed an EtOH diet for 6 weeks. The Comprehensive Lab Animal Monitoring System was used to measure metabolic phenotyping including CO_2_ production, O_2_ consumption, energy expenditure, and respiratory energy ratio. Data are presented as mean ± SEM (n = 5-9 per group). Each dot represents data from individual mice. A two-tailed unpaired *t*-test was used to compare WT and KO mice. \**P* < 0.05, \*\**P* < 0.01, \*\*\**P* < 0.001

### 3.2. Effects of CerS6 ablation on energy expenditure

We measured energy expenditure after 6 weeks on EtOH diet to evaluate the role of CerS6 in metabolic phenotypes. In male mice, CerS6 KO mice showed significantly higher O_2_ consumption, CO_2_ production, RER, and EE during the dark period but not during the light period (**Figure 1F-I**). In female mice, there were no differences in O_2_ consumption, CO_2_ production, RER, and EE between the EtOH-fed WT and CerS6 KO groups during dark and light periods (**Supplementary Figure 3A-D**). The RER (VCO_2_/ VO_2_) is an index of fuel oxidation. RER value of 0.7 indicates fat oxidation, and that of 1 indicates carbohydrate oxidation. Both male and female RER levels were above 0.7, suggesting that mice utilized a mix of fat and carbohydrate as a fuel source. Locomotor activity levels were similar between EtOH-fed WT and CerS6 KO mice of both sexes (**Supplementary Figure 3E, F**).

### 3.3. CerS6 ablation mitigates ethanol-induced glucose intolerance and insulin resistance

We previously reported that mice fed an EtOH diet developed glucose intolerance and insulin resistance as early as 4 weeks of feeding [5]. To determine the role of CerS6 deletion in insulin sensitivity, we performed GTT and ITT in EtOH-fed CerS6 KO and WT mice of both sexes. In both genders, CerS6 KO mice enhanced glucose tolerance during GTT as compared to WT animals (**Figure 2A, B**). During ITT, CerS6 KO led to a significant improvement in glucose clearance from circulation, indicating a strong insulin sensitizing effect compared to WT animals (**Figure 2C, D**).

**Figure 2.**
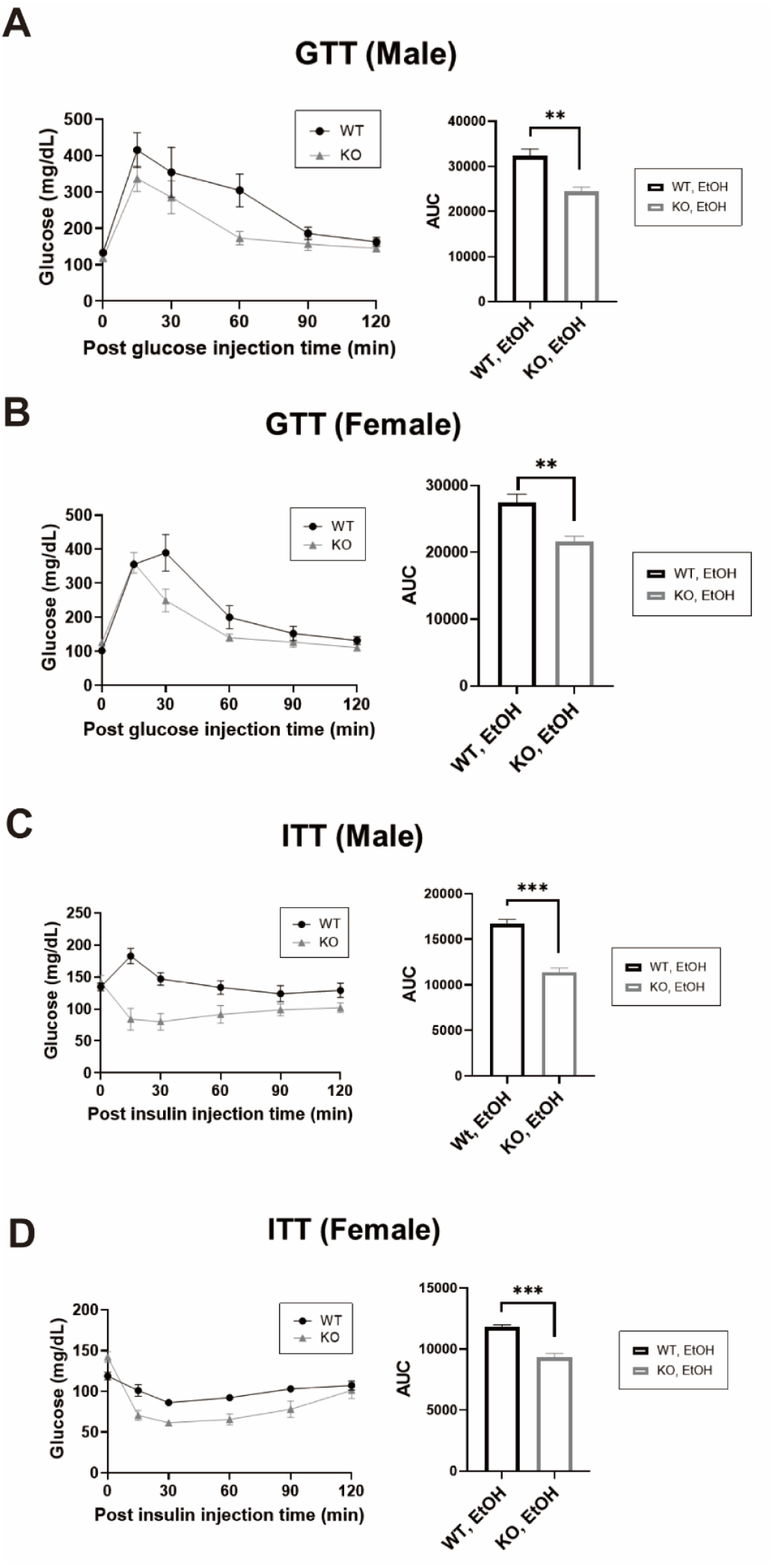
CerS6 ablation improves glucose intolerance and insulin sensitivity in the EtOH-fed mice. (A, B) Insulin tolerance test (ITT) and (C, D) glucose tolerance test (GTT) were performed after 6 weeks of feeding EtOH diet in WT and CerS6 KO mice of both sexes (n=5 per group). AUC, area under curve. Data are expressed as mean ± SEM. Each dot represents data from individual mice. A two-tailed unpaired *t*-test was used to compare WT and KO mice. \**P* < 0.05, \*\**P* < 0.01, \*\*\**P* < 0.001

### 3.4. CerS6 ablation attenuates ethanol-induced hepatic steatosis

EtOH diet promotes hepatic steatosis in a temporal pattern [5]. As expected from the body weight gain and dietary consumption, liver histology revealed that WT mice accumulate more hepatic lipids compared to the diet-matched KO mice (**Figure 3A**). There was no apparent evidence of inflammation or necrosis. In addition, there were no significant changes in hepatocellular injury between EtOH-fed WT and KO mice, as assessed by serum ALT (**Figure 3B**).

**Figure 3.**
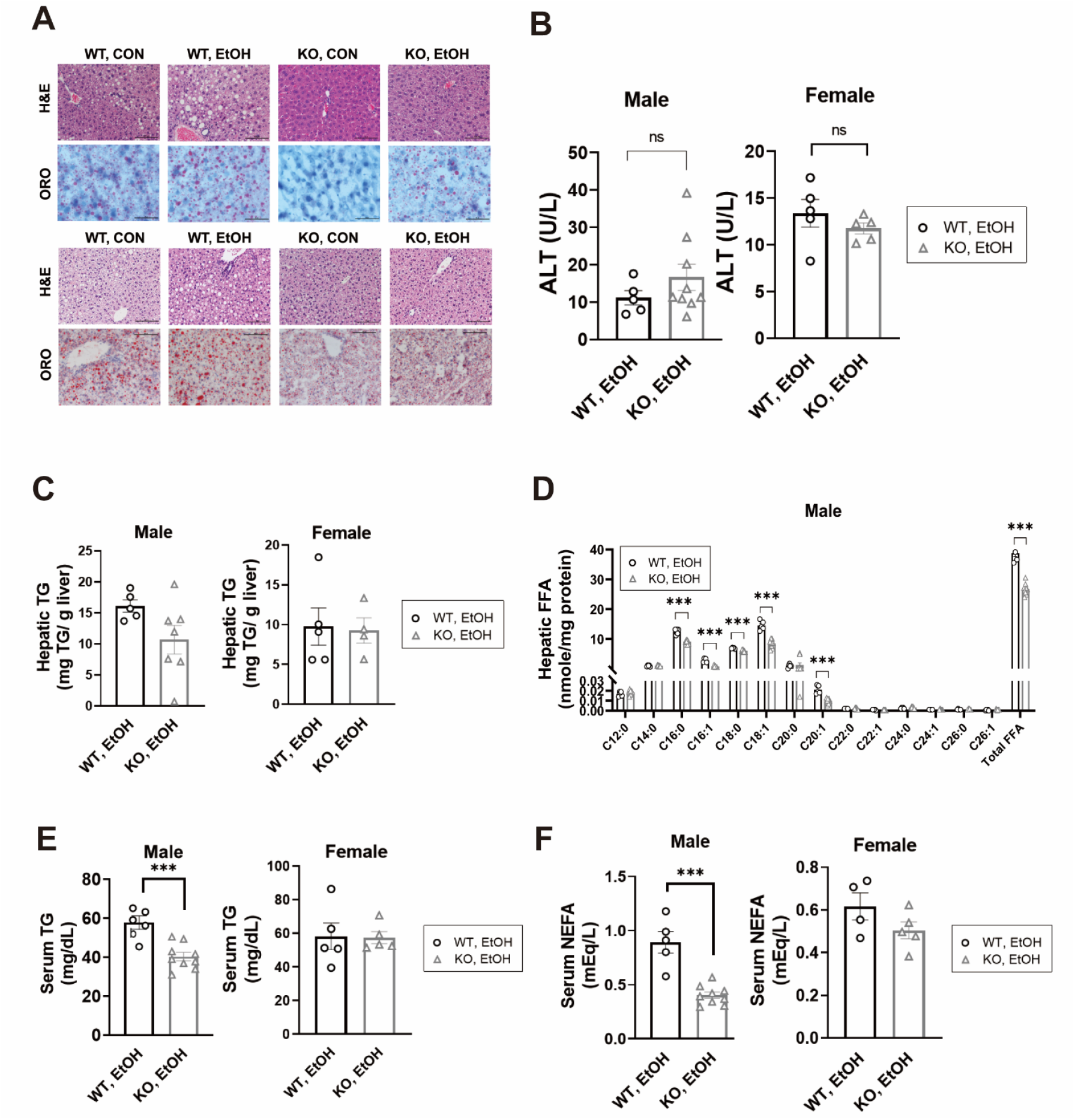
CerS6 ablation improves EtOH-induced hepatic steatosis and reduces serum lipids in the EtOH-fed mice. WT and CerS6 KO male and female mice were fed an EtOH diet for 6 weeks. **(A)** Representative histologic images of hematoxylin and eosin (H&E)-stained and Oil Red O (ORO)-stained liver sections. Original magnification ×20; n = 5-9 per group. **(B)** Serum ALT levels, **(C)** hepatic TG levels, **(D)** hepatic FFA levels, **(E)** serum TG levels, **(F)** serum NEFA levels of the EtOH-fed mice. Data are expressed as mean ± SEM. Each dot represents data from individual mice. A two-tailed unpaired *t*-test was used to compare WT and KO mice. \**P* < 0.05, \*\**P* < 0.01, \*\*\**P* < 0.001

We examined lipids that might explain the improvements in hepatic steatosis and measured triglycerides, NEFA, and sphingolipid levels in the liver and serum. In male mice, CerS6 deletion tended to decrease hepatic TGs (p=0.088), while there were no significant differences between WT and KO female mice (**Figure 3C**). Hepatic FFA levels were significantly decreased in EtOH-fed CerS6 KO mice compared to WT mice (**Figure 3D**). Among various species, long chain NEFA levels but not very long chain NEFA levels were significantly reduced in CerS6 KO male mice compared to WT mice. CerS6 deletion significantly reduced both circulating TG and NEFA levels in the EtOH-fed male mice but not in the EtOH-fed female mice (**Figure 3E, F**).

Sphingolipids in the liver and serum of EtOH-fed WT and KO mice were measured using mass spectrometry (**Table 1**). CerS6 deletion reduced C_16:0_-ceramide levels in both the serum and livers of both sexes, confirming a systemic effect of CerS6 deletion. Specifically, CerS6 deletion significantly reduced C_16:0_-ceramide in the liver by 29% and 83% in male and female mice, respectively; and in serum by 81% and 86% in male and female, respectively (**Figure 4A**). Similar to C_16:0_-ceramide, several long chain ceramides were significantly decreased in the liver and serum of KO mice of both sexes. In contrast, CerS6 deletion increased very long chain ceramides in the liver and serum of male mice. CerS6 deletion additionally reduced several species of dihydroceramides (a precursor of ceramide in the de novo ceramide synthetic pathway) in the livers and serum of EtOH-fed mice (**Figure 4B, C**). We also calculated serum ceramide ratios comparing the relative proportion of very long and long-chain ceramides (C_24:0_/C_16:0_) (**Supplementary Table 1**). The ceramide ratio has been suggested as a new serum marker for metabolic dysfunction based on various observations about the strong association between the ceramide ratio and insulin resistance as well as cardiovascular outcomes [19]. C_24:0_/C_16:0_ ratio was significantly increased by CerS6 deletion in the serum of EtOH-fed male and female mice. Hepatic levels of dihydrosphingosine (dhSph), sphingosine (Sph) and their phosphorylated derivatives were not altered by CerS6 ablation in either female or male EtOH-fed mice (**Supplementary Figure 4**). CerS6 significantly decreased dhSph-1P and Sph-1P levels in the serum of female mice but not in that of male mice (**Figure 4D**).

**Table 1.**
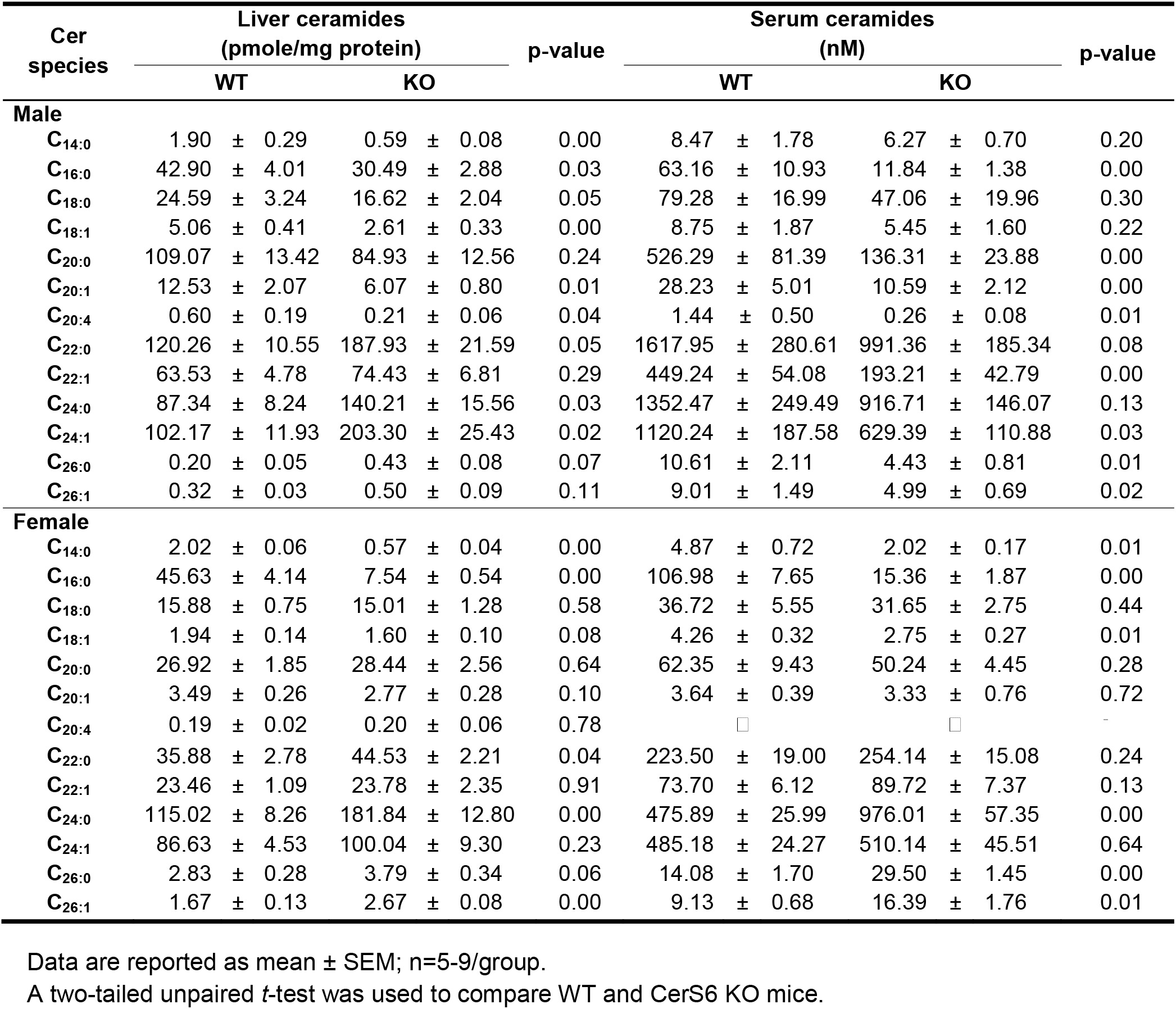
Hepatic and serum ceramide content of ethanol (EtOH)-fed ceramide synthase 6 (CerS6) knockout and wild-type mice

**Figure 4.**
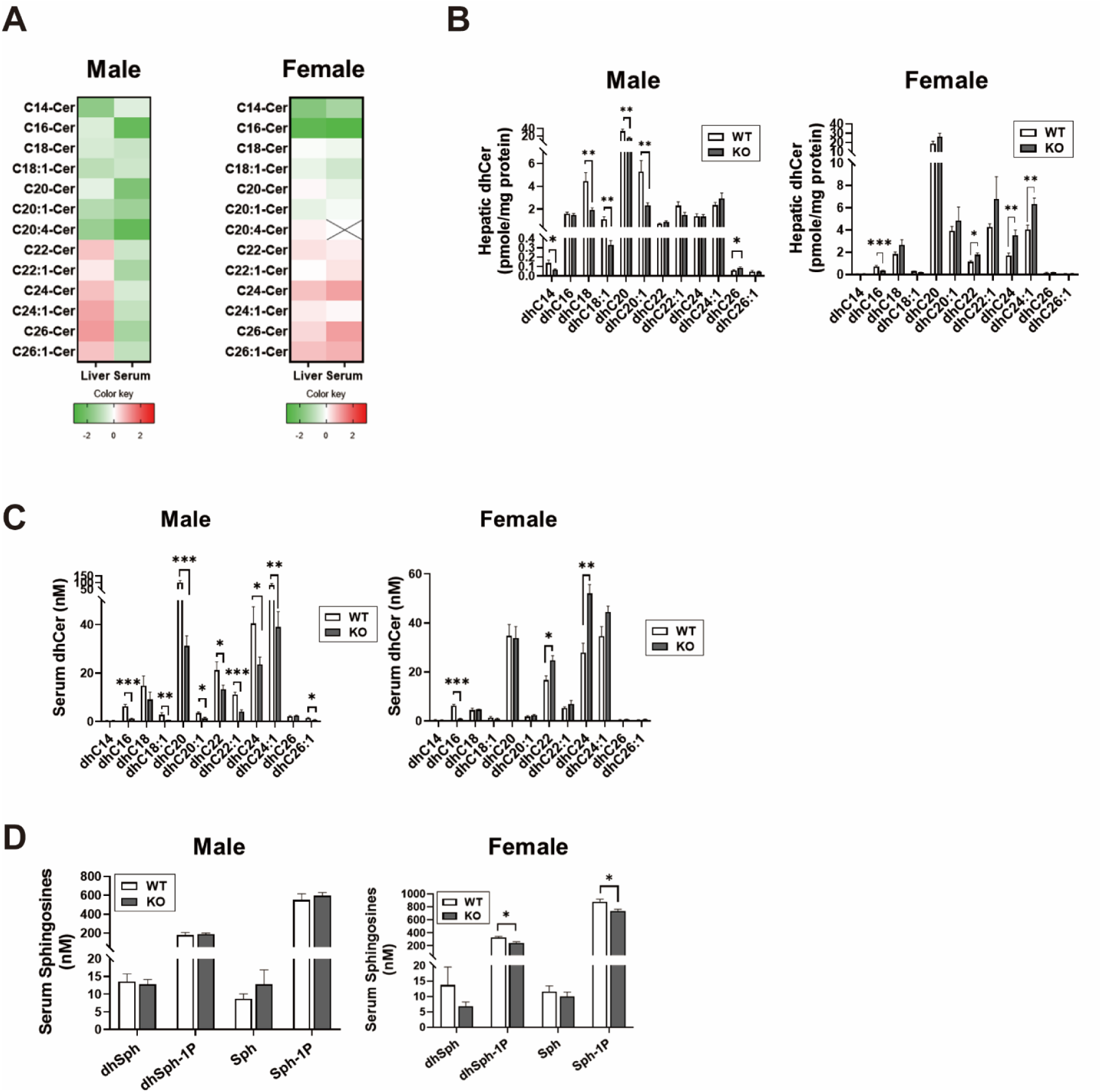
CerS6 ablation alters sphingolipids in the liver and serum of EtOH-fed WT and CerS6 KO mice. WT and CerS6 KO male and female mice were fed an EtOH diet for 6 weeks. **(A)** In the heatmap, the color code indicates the log2 of the ratio between means of the groups for an individual ceramide. The x axes denote the ceramide species. A more intense red color indicates a greater increase of absolute concentration of the individual ceramide in the liver and serum. **(B, C)** Dihydroceramide (dhCer), **(D)** dihydrosphingosine (dhSph), dihydrosphingosine-1-phosphate (dhSph-1P), sphingosine (Sph), and sphingosine-1-phosphate (Sph-1P) levels in the livers and serum of WT and KO mice. Data are expressed as means ± SEM (n=5). A two-tailed unpaired *t*-test was used to compare WT and KO mice. \**P* < 0.05, \*\**P* < 0.01, \*\*\**P* < 0.001

### 3.5. RNA-seq revealed differentially expressed genes and pathways by EtOH diet and CerS6 ablation

To gain insight into the molecular mechanisms of the role of CerS6 and delineate the differentially expressed genes (DEGs) in CerS6 KO compared to WT mice, we performed RNA-seq in the liver samples from CON- or EtOH-fed WT and CerS6 KO male mice. In the principal component analysis, CON-fed WT and KO mice revealed a clear separation and EtOH-fed WT and KO mice showed a slight separation of liver transcriptomes (**Figure 5A**). A heatmap depicts > 2-fold changed DEGs with the false discovery rate, P < 0.05 in CON-fed or EtOH-fed KO mice and WT mice. Volcano plots show that 74 genes are significantly upregulated, and 66 genes are downregulated by CerS6 deletion in EtOH-fed male mice (**Figure 5C**). Among these genes, gene enrichment analysis identified several significantly altered metabolic pathways in EtOH-fed KO versus EtOH-fed WT mice, including organic acid biosynthetic process, triglyceride biosynthetic process, positive regulation of lipid storage, inflammatory response, and fat cell differentiation (**Figure 5D**). In addition, KEGG pathways associated with lipid metabolism were also identified in both genotype and EtOH diet effects.

**Figure 5.**
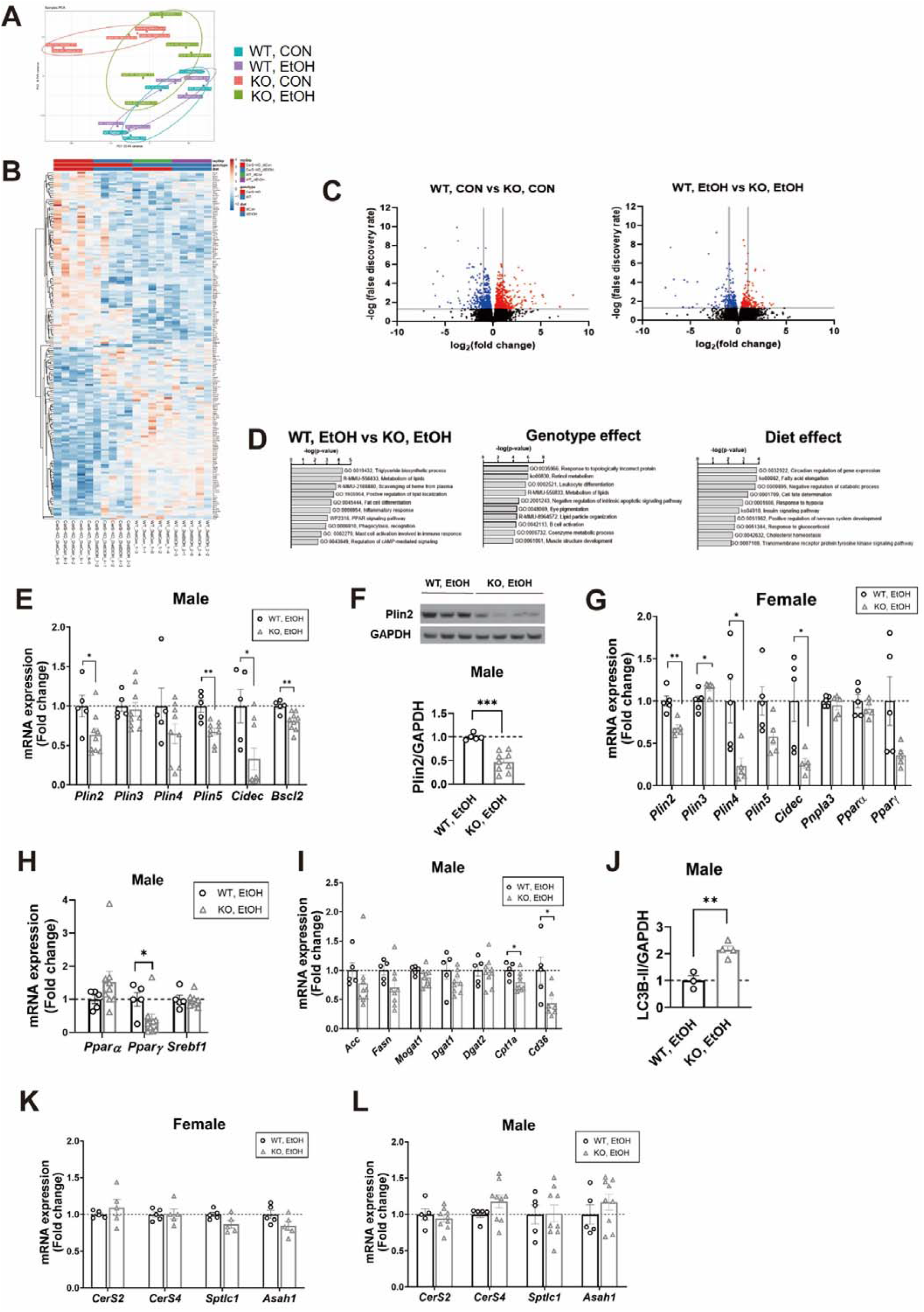
RNA-seq analysis of liver transcriptomes in male WT and CerS6 KO mice fed a CON or an EtOH diet for six weeks. **(A)** Principal component analysis (PCA) showing global sample distribution profiles. Each dot represents data from individual mice. (**B**) Heatmap analysis of the differentially expressed genes regulated by genotype and diet in four groups. **(C)** Volcano plot of the DEGs in the liver of diet-matched WT vs. KO mice. Upregulated genes are in red and downregulated genes are in blue. Vertical lines indicate the threshold for a relative expression fold change of > 2 or < -2 compared to WT and horizontal line indicates the threshold of false discovery rate < 0.05. **(D)** Top 10 pathways based on the pathway enrichment analysis results of the genes that are significantly regulated in EtOH-fed KO mice compared to EtOH-fed WT mice, and that are significantly regulated by diet and genotype in four groups. **(E-L)** CerS6 deletion alters gene expression related to lipid metabolism in the livers of EtOH-fed mice. Relative mRNA expression levels were determined by real-time RT-PCR and were normalized to 36B4 mRNA levels. Relative protein levels by Western Blotting were normalized to GAPDH. Data are expressed as means ± SEM (n=5/group). A two-tailed unpaired *t*-test was used to compare WT and KO mice. \**P* < 0.05, \*\**P* < 0.01, \*\*\**P* < 0.001

### 3.6. CerS6 ablation reduces hepatic expression of lipid droplet-associated proteins

Previously, EtOH diet was shown to promote hepatic steatosis with *Plin2* upregulation, a major lipid droplet-associated protein [5]. CerS6 deletion significantly decreased protein and mRNA expression of PLIN2 in the livers of EtOH-fed mice (**Figure 5E-G**). Based on the RNA-seq data, we also measured gene expression regarding lipid droplet biology and demonstrated that CerS6 ablation significantly decreased mRNA expression of *Plin5*, cell death-inducing DNA fragmentation factor A– like effector C (*Cidec*), and Berardinelli-Seip congenital lipodystrophy (*Bscl2*) in the livers of EtOH-fed male mice (**Figure 5E**). Similarly, CerS6 deletion significantly decreased mRNA expression of *Plin4* and *Cidec* in the livers of EtOH-fed female mice (**Figure 5G**). CerS6 deletion significantly reduced peroxisome proliferator-activated receptor γ (*Pparg)* and *Cd36* in male mice and tended to decrease *Pparg* mRNA expression in female mice (**Figure 5G-I**). In addition, the autophagosome marker LC3-II protein levels in CerS6 KO mice were significantly increased compared with WT mice (**Figure 5J**). CerS6 deletion didn’t impact other ceramide synthetic genes such as *CerS2, CerS4, Spt*, or *Asah1* in EtOH-fed male and female mice, confirming that there was no compensatory effect of CerS6 deletion in sphingolipid metabolism (**Figure 5K, L**).

### 3.7. In vitro studies

Previously, it has been demonstrated that PLIN2 is regulated by CerS6 in ALD pathogenesis [10], also confirmed in the current in vivo study. To further elucidate the role of CerS6 and CerS6-derived C_16:0_-ceramide in PLIN2 regulation in a human hepatocyte cell-line, CerS6 null VL17A cells were generated by CRISPR/Cas9. In response to 100mM EtOH, PLIN2 protein levels were upregulated compared to control in parental VL17A cells, but not in CerS6 KO cells (**Figure 6A**). C_16:0_-ceramide were treated in parental and CerS6 KO VL17A cells. 100nM of C_16:0_-ceramide was not cytotoxic in hepatocytes, as determined by MTT assays (data not shown) and these concentrations were similar to circulating levels found in EtOH-fed WT mice. Co-treatment of C_16:0_-ceramide and ethanol increased PLIN2 in parental cells but not in CerS6 KO cells, indicating that CerS6 but not C_16:0_-ceramide per se is associated with increased PLIN2 protein expression (**Figure 6B**).

**Figure 6.**
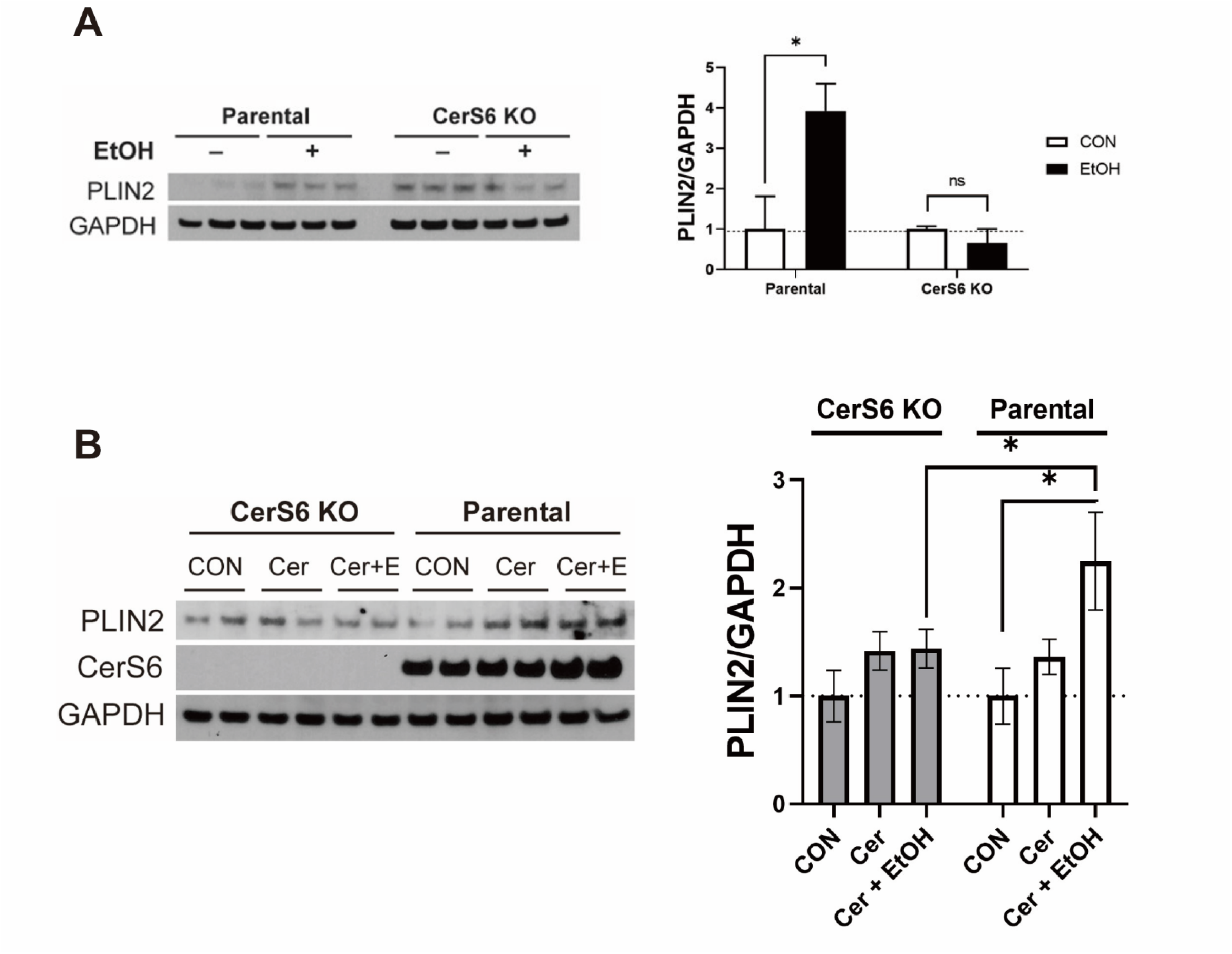
CerS6 ablation alters perilipin2 (PLIN2) expression in human hepatocytes. **(A)** PLIN2 expression in cell lysates of VL17A parental and CerS6 KO cells that were treated with control or 100mM ethanol for 48hr (n=3); **(B)** with control, 100nM C_16:0_-ceramide, or 100nM C_16:0_-ceramide with 100mM ethanol for 24hr (n=4). Data are expressed as mean ± SEM. Statistical analyses were performed using a two-tailed unpaired *t*-test or one-way ANOVA with Tukey’s posthoc test. \**P* < 0.05.

### 3.8. Hepatic CerS6 expression in patients with liver diseases

We previously demonstrated that CERS6 is upregulated in the livers of patients with alcoholic steatosis, as determined by immunohistochemistry [10]. Here, we analyzed hepatic RNAseq profiling from patients with different stages of ALD as well as diseased controls. Patients with more severe forms of ALD (i.e. severe acute alcohol hepatitis) had the highest levels of hepatic *CERS6* mRNA expression, followed by patients with non-severe acute alcohol hepatitis (**Figure 7**). These results demonstrate that not only is CerS6 upregulation an early biomarker of ALD but increases with disease progression. In addition to *CERS6, PLIN3* and *CIDEC* mRNA expression increased with disease progression.

**Figure 7.**
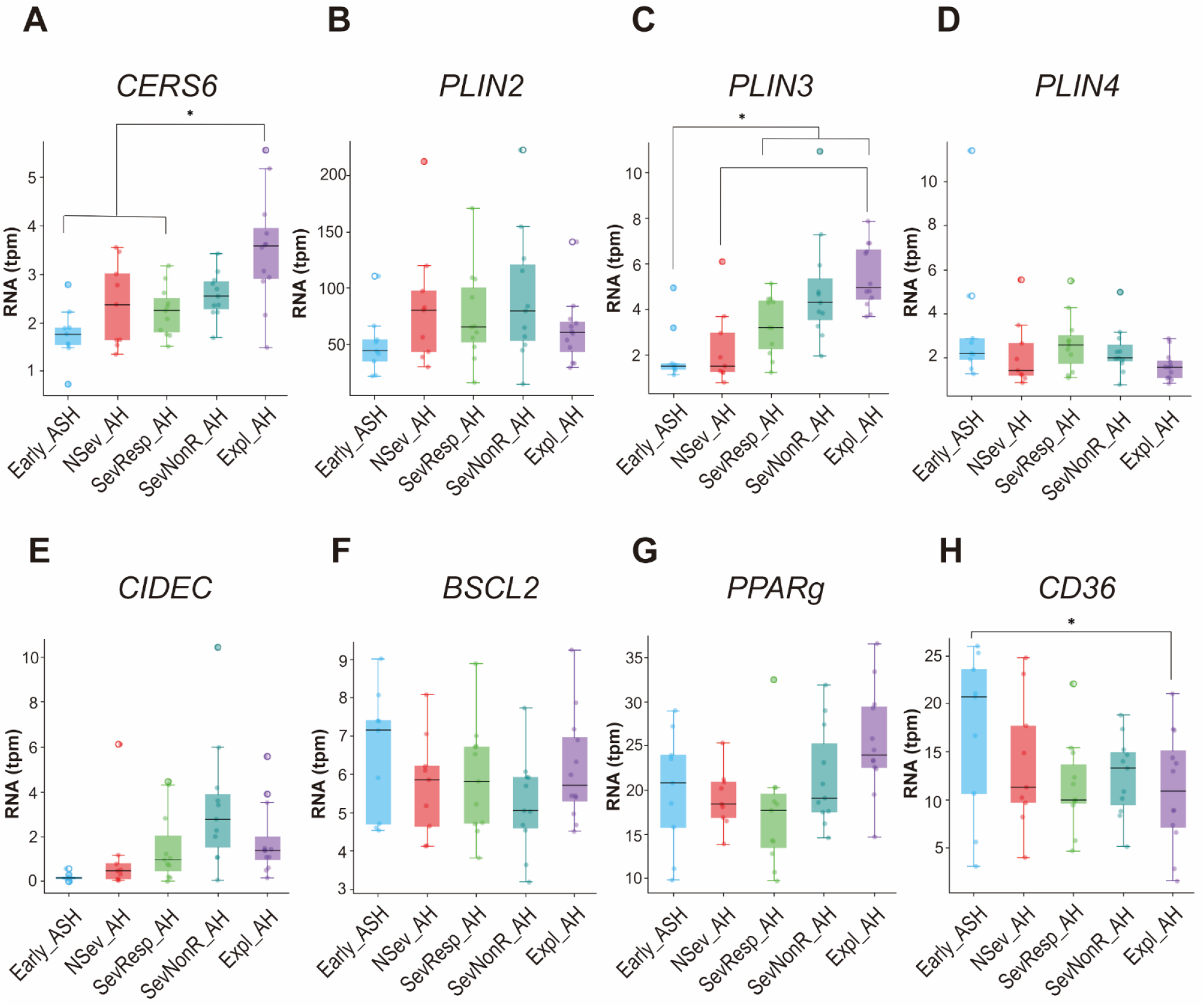
Hepatic gene expression in human subjects with different stages of ALD and diseased controls. Early ASH, early acute alcoholic hepatitis (*N* = 12); Nsev_AH, nonsevere acute alcoholic hepatitis (*N* = 11); SevResp_AH, severe alcoholic hepatitis responders to medical therapy (*N* = 9); SevNonR_AH, severe alcoholic hepatitis non-responders to medical therapy (*N* = 9); Expl_AH, explants from patients with alcoholic hepatitis (*N* = 11). Gene expression levels are presented in transcripts per million reads (tpm). Data are expressed as mean ± SEM. Statistical analyses were performed using a two-tailed unpaired *t*-test or one-way ANOVA with Tukey’s posthoc test. \**P* < 0.05.

## 4. Discussion

Excessive ceramide accumulation is known to be a key pathological feature of ALD in both human patients and animal models [11]. CerS6, the enzyme involved in ceramide *de novo* synthesis and salvage pathways, and CerS6-derived C_16:0_-ceramides are emerging as major contributors in the development of various metabolic diseases including insulin resistance, obesity and NAFLD [20]. However, the role of CerS6 in ALD is incompletely understood. We and others have shown that overall ceramide reduction by using pharmacological and genetic manipulations can alleviate EtOH-induced fatty liver [16, 21, 22]. In this study, we explore the role of CerS6 in alleviating EtOH-induced hepatic steatosis and insulin resistance, using WT and KO mice with both sexes and human hepatoma cells.

Here, we revealed that CerS6 deletion protected against body weight gain in both sexes. Similarly, studies have shown that conventional or conditional CerS6 deletion alleviated diet-induced obesity in in vivo models [9, 23]. CerS6 knockdown using antisense oligonucleotides also exhibited its protective effect against weight gain and insulin-resistance [24]. We speculate that decreased body weight gain in CerS6 KO mice might be due to reduced food consumption and an increased rate of energy expenditure, which agrees with previous studies. Accrual of hypothalamic ceramides stimulates orexigenic pathways, dysregulates energy balance, and further perturbs glucose homeostasis in mice models [25]. In addition, central administration of a cell-permeable C_6:0_-ceramide analogue to the hypothalamus increases hypothalamic C_16:0_-ceramides, and subsequently increases body weight, which might emphasize the pathological role of C_16:0_-ceramides in obesity [26]. In this regard, our study corroborates the CerS6’s role as a metabolic regulator in ALD animal models. Our data indicates that CerS6 deletion alleviated insulin resistance and glucose intolerance in both sexes. The role of ceramides in insulin resistance has been established in ALD development [27]. Specifically, CerS6-derived C_16:0_-ceramides have been demonstrated to impair insulin action in liver and brown adipose tissue [9]. One of the underlying mechanisms in ALD is that EtOH-induced insulin resistance exacerbates adipose tissue lipolysis and then increases circulating NEFAs [28]. Indeed, we have shown that alcohol increases serum NEFAs, and CerS6 deletion downregulates fatty acid translocase CD36 that participates in hepatic uptake of fatty acids [29]. This is in line with previous studies that ceramides induce CD36 translocation and stimulate fatty acid uptake in murine livers [30, 31].

Notably, we demonstrated that CerS6 regulates lipid droplet biogenesis. CerS6 deletion down-regulates a major lipid droplet-associated protein, PLIN2. We have previously reported a temporal relationship between PLIN2 up-regulation and hepatic ceramide accumulation in *in vivo* ALD models [5]. In our *in vitro* model of alcoholic steatosis, using human hepatoma VL17A cells, we demonstrated that CerS6 upregulates PLIN2 expression in response to EtOH, which is not mediated by C_16:0_-ceramides. This is supported by the recent finding that CerS has a dual role as an enzyme that produces ceramides as well as a transcription regulator [17]. In addition to PLIN2, we also demonstrated that other LD associated proteins were downregulated in CerS6 KO mice, suggesting that CerS6 deletion could efficiently blunt lipid droplet formation in the early stage of ALD. Interestingly, we exhibited that ceramide profiles in serum and liver differ according to sex. Sex-dependent ceramide levels were reported in various tissues including heart [32], kidney [33], cerebral cortex [34], and plasma [35]. These differential composition of ceramides might be explained by the action of estradiol [36]. Despite the sex-dependent ceramide profiles, we confirm that CerS6 deletion could alleviate hepatic steatosis in both sexes.

Finally, our results have implications for human ALD. Previously, we reported that CerS6 is up-regulated in the livers of human alcoholic patients and alcohol-fed mice as well as in the lipid droplet fractions of EtOH-treated VL17A cells [10]. We now add to this growing body of literature and demonstrate the association between CerS6 expression and disease severity. These findings reveal that CerS6-derived ceramides could be an attractive therapeutic target for treating ALD. Additionally, we found that several lipid droplet-associated proteins were also associated with disease progression, showing the similar pattern in mice.

One limitation to the present study was the use of conventional knockout animals, which allowed us to reveal the role of CerS6 in whole body metabolism. As CerS6-derived C_16:0_ ceramides can act at multiple levels including adipose tissue, liver, brain, we are unable to establish here the tissue-specific contributions of CerS6 in ALD pathogenesis. The tissue-specific effects of CerS6 in ALD are of interest for future studies.

In conclusion, our findings demonstrate that CerS6 deletion attenuates EtOH-induced hepatic steatosis and alleviates lipid and glucose metabolic dysfunction in mice fed an EtOH diet. Based on untargeted gene expression analysis, the mechanism does appear to be from reduced lipid droplet biogenesis.

## List of Abbreviations

ALD: alcohol-associated liver disease
ALT: alanine aminotransferase
Bscl: Berardinelli-Seip congenital lipodystrophy
CerS: ceramide synthase
Cidec: cell death-inducing DNA fragmentation factor A–like effector C
CON: control
dhSph: dihydrosphingosine
DEGs: differentially expressed genes
eWAT: epididymal white adipose tissue
EE: energy expenditure
EtOH: ethanol
GTT: glucose tolerance test
H&E: hematoxylin and eosin
ITT: insulin tolerance test
KO: knockout
LD: lipid droplet
NEFAs: non-esterified FAs
ORO: Oil-red-O
Plin: perilipin
PPAR: peroxisome proliferator-activated receptor
RER: respiratory exchange ratio
SEM: standard error of measurement
SPT: serine palmitoyl transferase
Sph: sphingosine
TBST: tris-buffered saline Tween 20
TGs: triglycerides
WT: wild-type

## 5. Acknowledgement

This study was supported by NIH grant (1R01AA026302-01, P30 DK050306). The authors would like to thank the Molecular Pathology and Imaging Core (NIH-P30-DK050306), Dr. Jonathan Schug, and Dr. John Tobias for expert technical, and Dr. Dahn Clemens for the generous gift of VL17A cells. The lipidomics studies were performed at the Lipidomics Core Facility at the Medical University of South Carolina (MUSC), supported by NIH (P30 CA138313 and P30 GM103339).

## References

[1] Asrani, S.K., Mellinger, J., Arab, J.P., Shah, V.H., 2021. Reducing the Global Burden of Alcohol-Associated Liver Disease: A Blueprint for Action. Hepatology 73(5):2039–2050.

[2] Andersen, B.N., Hagen, C., Faber, O.K., Lindholm, J., Boisen, P., Worning, H., 1983. Glucose tolerance and B cell function in chronic alcoholism: Its relation to hepatic histology and exocrine pancreatic function. Metabolism: Clinical and Experimental 32(11):1029–1032.

[3] Scorletti, E., Carr, R.M., 2021. A new perspective on NAFLD: focusing on lipid droplets. Journal of Hepatology.

[4] Whitfield, J.B., Schwantes-An, T.-H., Darlay, R., Aithal, G.P., Atkinson, S.R., Bataller, R., et al., 2022. A genetic risk score and diabetes predict development of alcohol-related cirrhosis in drinkers. Journal of Hepatology 76(2):275–282.

[5] Carr, R.M., Dhir, R., Yin, X., Agarwal, B., Ahima, R.S., 2013. Temporal effects of ethanol consumption on energy homeostasis, hepatic steatosis, and insulin sensitivity in mice. Alcoholism, Clinical and Experimental Research 37(7):1091–1099.

[6] Carr, R.M., Peralta, G., Yin, X., Ahima, R.S., 2014. Absence of perilipin 2 prevents hepatic steatosis, glucose intolerance and ceramide accumulation in alcohol-fed mice. PloS One 9(5):e97118.

[7] Mullen, Thomas D., Hannun Yusuf A., Obeid Lina M., 2012. Ceramide synthases at the centre of sphingolipid metabolism and biology. Biochemical Journal 441(3):789–802.

[8] Mizutani, Y., Kihara, A., Igarashi, Y., 2005. Mammalian Lass6 and its related family members regulate synthesis of specific ceramides. Biochemical Journal 390(1):263–271.

[9] Turpin, Sarah M., Nicholls Hayley T., Willmes Diana M., Mourier, A., Brodesser, S., Wunderlich Claudia M., et al., 2014. Obesity-Induced CerS6-Dependent C16:0 Ceramide Production Promotes Weight Gain and Glucose Intolerance. Cell Metabolism 20(4):678–686.

[10] Williams, B., Correnti, J., Oranu, A., Lin, A., Scott, V., Annoh, M., et al., 2018. A novel role for ceramide synthase 6 in mouse and human alcoholic steatosis. FASEB Journal 32(1):130–142.

[11] Jeon, S., Carr, R., 2020. Alcohol effects on hepatic lipid metabolism. Journal of Lipid Research 61(4):470–479.

[12] Holland, W.L., Summers, S.A., 2018. Strong Heart, Low Ceramides. Diabetes 67(8):1457–1460.

[13] Longato, L., Ripp, K., Setshedi, M., Dostalek, M., Akhlaghi, F., Branda, M., et al., 2012. Insulin resistance, ceramide accumulation, and endoplasmic reticulum stress in human chronic alcohol-related liver disease. Oxidative Medicine and Cellular Longevity 2012.

[14] Liangpunsakul, S., Rahmini, Y., Ross, R.A., Zhao, Z., Xu, Y., Crabb, D.W., 2011. Imipramine blocks ethanol-induced ASMase activation, ceramide generation, and PP2A activation, and ameliorates hepatic steatosis in ethanol-fed mice. American Journal of Physiology-Gastrointestinal and Liver Physiology 302(5):G515–G523.

[15] Tong, M., Longato, L., Ramirez, T., Zabala, V., Wands, J.R., de la Monte, S.M., 2014. Therapeutic reversal of chronic alcohol-related steatohepatitis with the ceramide inhibitor myriocin. International Journal of Experimental Pathology 95(1):49–63.

[16] Correnti, J., Lin, C., Brettschneider, J., Kuriakose, A., Jeon, S., Scorletti, E., et al., 2020. Liver-specific ceramide reduction alleviates steatosis and insulin resistance in alcohol-fed mice. Journal of Lipid Research 61(7):983–994.

[17] Sociale, M., Wulf, A.-L., Breiden, B., Klee, K., Thielisch, M., Eckardt, F., et al., 2018. Ceramide Synthase Schlank Is a Transcriptional Regulator Adapting Gene Expression to Energy Requirements. Cell Reports 22(4):967–978.

[18] Argemi, J., Latasa, M.U., Atkinson, S.R., Blokhin, I.O., Massey, V., Gue, J.P., et al., 2019. Defective HNF4alpha-dependent gene expression as a driver of hepatocellular failure in alcoholic hepatitis. Nature Communications 10(1):3126.

[19] Tippetts, T.S., Holland, W.L., Summers, S.A., 2018. The ceramide ratio: a predictor of cardiometabolic risk1. Journal of Lipid Research 59(9):1549–1550.

[20] Turpin-Nolan, S.M., Brüning, J.C., 2020. The role of ceramides in metabolic disorders: when size and localization matters. Nature Reviews: Endocrinology 16(4):224–233.

[21] Correnti, J.M., Juskeviciute, E., Swarup, A., Hoek, J.B., 2014. Pharmacological ceramide reduction alleviates alcohol-induced steatosis and hepatomegaly in adiponectin knockout mice. American Journal of Physiology-Gastrointestinal and Liver Physiology 306(11):G959–G973.

[22] Lizarazo, D., Zabala, V., Tong, M., Longato, L., de la Monte, S.M., 2013. Ceramide inhibitor myriocin restores insulin/insulin growth factor signaling for liver remodeling in experimental alcohol-related steatohepatitis. Journal of Gastroenterology and Hepatology 28(10):1660–1668.

[23] Hammerschmidt, P., Ostkotte, D., Nolte, H., Gerl, M.J., Jais, A., Brunner, H.L., et al., 2019. CerS6-Derived Sphingolipids Interact with Mff and Promote Mitochondrial Fragmentation in Obesity. Cell 177(6):1536-1552.e1523.

[24] Raichur, S., Brunner, B., Bielohuby, M., Hansen, G., Pfenninger, A., Wang, B., et al., 2019. The role of C16:0 ceramide in the development of obesity and type 2 diabetes: CerS6 inhibition as a novel therapeutic approach. Molecular Metabolism 21:36–50.

[25] Magnan, C., Le Stunff, H., 2021. Role of hypothalamic de novo ceramides synthesis in obesity and associated metabolic disorders. Molecular Metabolism:101298.

[26] Contreras, C., González-García, I., Martínez-Sánchez, N., Seoane-Collazo, P., Jacas, J., Morgan Donald A., et al., 2014. Central Ceramide-Induced Hypothalamic Lipotoxicity and ER Stress Regulate Energy Balance. Cell Reports 9(1):366–377.

[27] Carr, R.M., Correnti, J., 2015. Insulin resistance in clinical and experimental alcoholic liver disease. Annals of the New York Academy of Sciences 1353:1–20.

[28] Wei, X., Shi, X., Zhong, W., Zhao, Y., Tang, Y., Sun, W., et al., 2013. Chronic alcohol exposure disturbs lipid homeostasis at the adipose tissue-liver axis in mice: analysis of triacylglycerols using high-resolution mass spectrometry in combination with in vivo metabolite deuterium labeling. PloS One 8(2):e55382.

[29] Clugston, R.D., Yuen, J.J., Hu, Y., Abumrad, N.A., Berk, P.D., Goldberg, I.J., et al., 2014. CD36-deficient mice are resistant to alcohol- and high-carbohydrate-induced hepatic steatosis. Journal of Lipid Research 55(2):239–246.

[30] Chaurasia, B., Tippetts, T.S., Mayoral Monibas, R., Liu, J., Li, Y., Wang, L., et al., 2019. Targeting a ceramide double bond improves insulin resistance and hepatic steatosis. Science 365(6451):386–392.

[31] Xia, Jonathan Y., Holland William L., Kusminski Christine M., Sun, K., Sharma Ankit X., Pearson Mackenzie J., et al., 2015. Targeted Induction of Ceramide Degradation Leads to Improved Systemic Metabolism and Reduced Hepatic Steatosis. Cell Metabolism 22(2):266–278.

[32] Kadokami, T., McTiernan, C.F., Kubota, T., Frye, C.S., Feldman, A.M., 2000. Sex-related survival differences in murine cardiomyopathy are associated with differences in TNF-receptor expression. The Journal of clinical investigation 106(4):589–597.

[33] Durant, B., Forni, S., Sweetman, L., Brignol, N., Meng, X.-L., Benjamin, E.R., et al., 2011. Sex differences of urinary and kidney globotriaosylceramide and lyso-globotriaosylceramide in Fabry mice. Journal of Lipid Research 52(9):1742–1746.

[34] Barrier, L., Ingrand, S., Fauconneau, B., Page, G., 2010. Gender-dependent accumulation of ceramides in the cerebral cortex of the APPSL/PS1Ki mouse model of Alzheimer’s disease. Neurobiology of Aging 31(11):1843–1853.

[35] Weir, J.M., Wong, G., Barlow, C.K., Greeve, M.A., Kowalczyk, A., Almasy, L., et al., 2013. Plasma lipid profiling in a large population-based cohort. Journal of Lipid Research 54(10):2898–2908.

[36] González-García, I., Contreras, C., Estévez-Salguero, Á., Ruíz-Pino, F., Colsh, B., Pensado, I., et al., 2018. Estradiol Regulates Energy Balance by Ameliorating Hypothalamic Ceramide-Induced ER Stress. Cell Reports 25(2):413-423.e415.

