## Supplemental data for "Ceramide synthase 6 (CerS6) promotes alcohol-induced fatty liver by promoting metabolic dysfunction and upregulating lipid droplet-associated proteins"

**Supplementary Data**

**Supplementary Figure 1**. Confirmation of ceramide synthase (CerS6) KO mice. **(A)** Genotyping of WT, heterozygous, and CerS6 KO mice using tail DNA. Bands correspond to 295bp KO PCR product and 460 bp WT PCR product. **(B)** Immunoblots of CerS6 and GAPDH from liver and epididymal white adipose tissue (eWAT) lysates of WT and CerS6 KO mice.

**(A)**

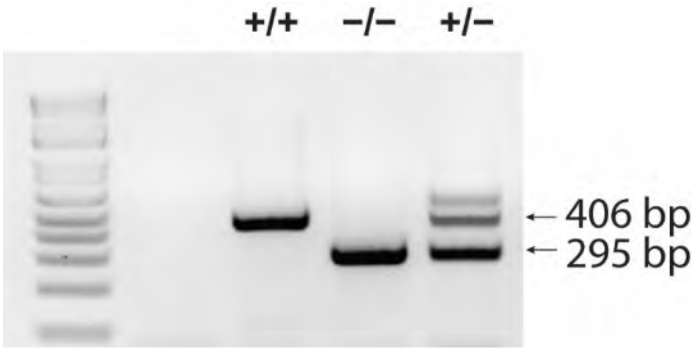


**(B)**

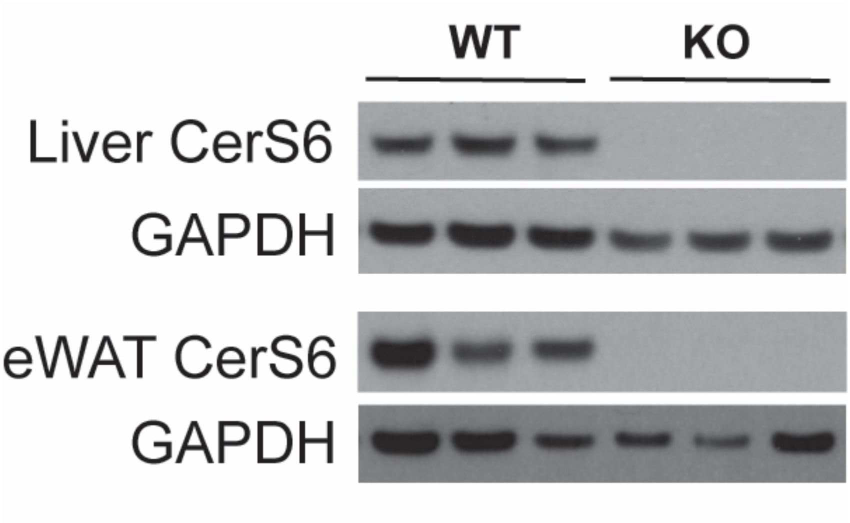

**Supplementary Figure 2**. Body weight changes over the feeding period in the CON-fed or EtOH-fed WT and CerS6 KO male and female mice. WT and CerS6 KO male and female mice were fed an EtOH diet for 6 weeks. Data are presented as mean ± SEM (n = 5-9 per group). A two-tailed unpaired *t*-test was used to compare WT and KO mice. EDC: ethanol derived calories, ns: not significant

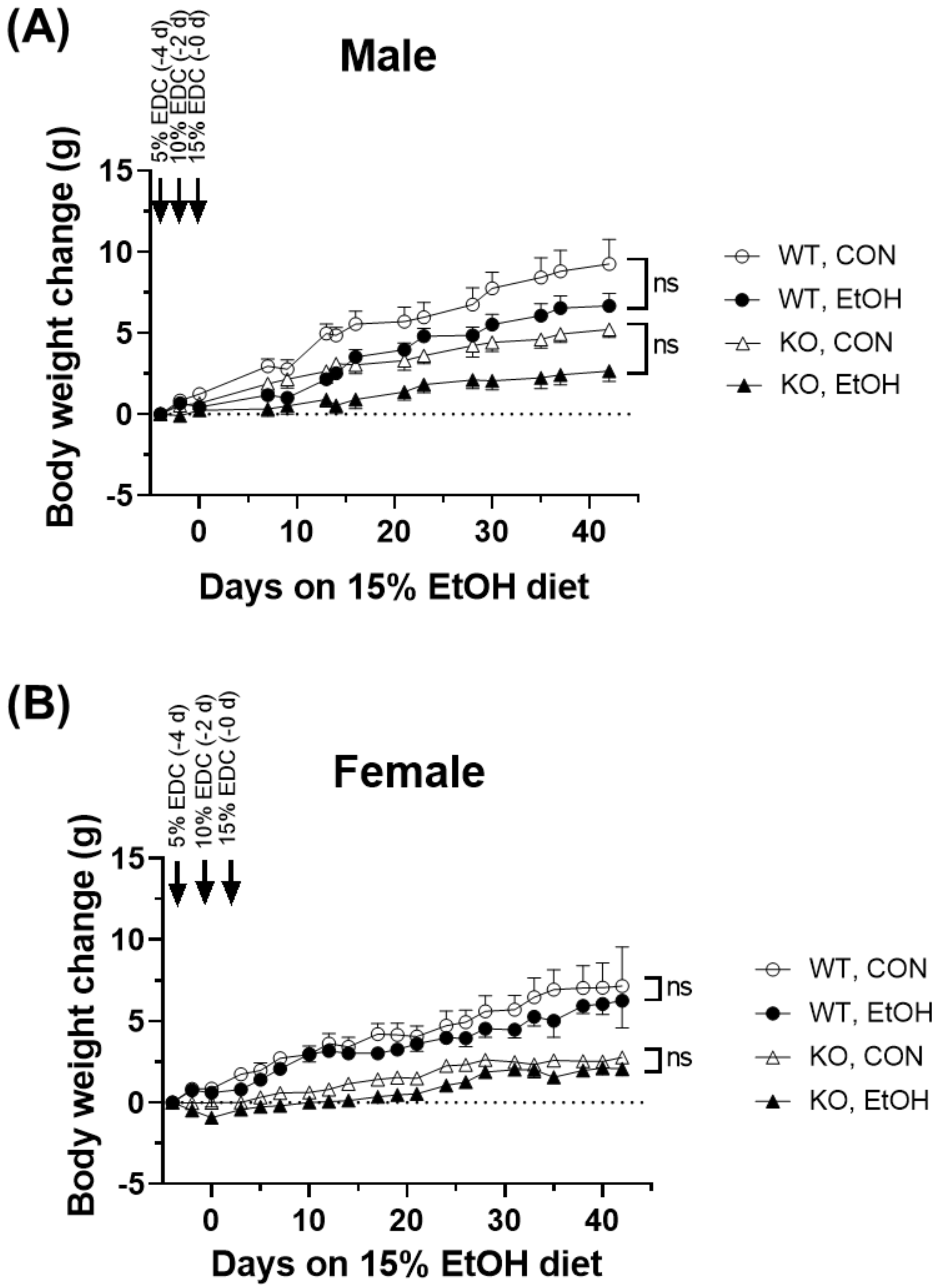

**Supplementary Figure 3.** CerS6 ablation has no effect on **(A-D)** metabolic phenotyping in EtOH-fed female mice, and **(E, F)** locomotor activity in EtOH-fed male and female mice. WT and CerS6 KO male and female mice were fed an EtOH diet for 6 weeks. The Comprehensive Lab Animal Monitoring System was used to measure metabolic phenotyping including $CO_2$ production, $O_2$ consumption, energy expenditure, and respiratory energy ratio. Data are expressed as mean ± SEM (n=5). A two-tailed unpaired *t*-test was used to compare WT and KO mice. **P* < 0.05, ***P* < 0.01, ****P* < 0.001

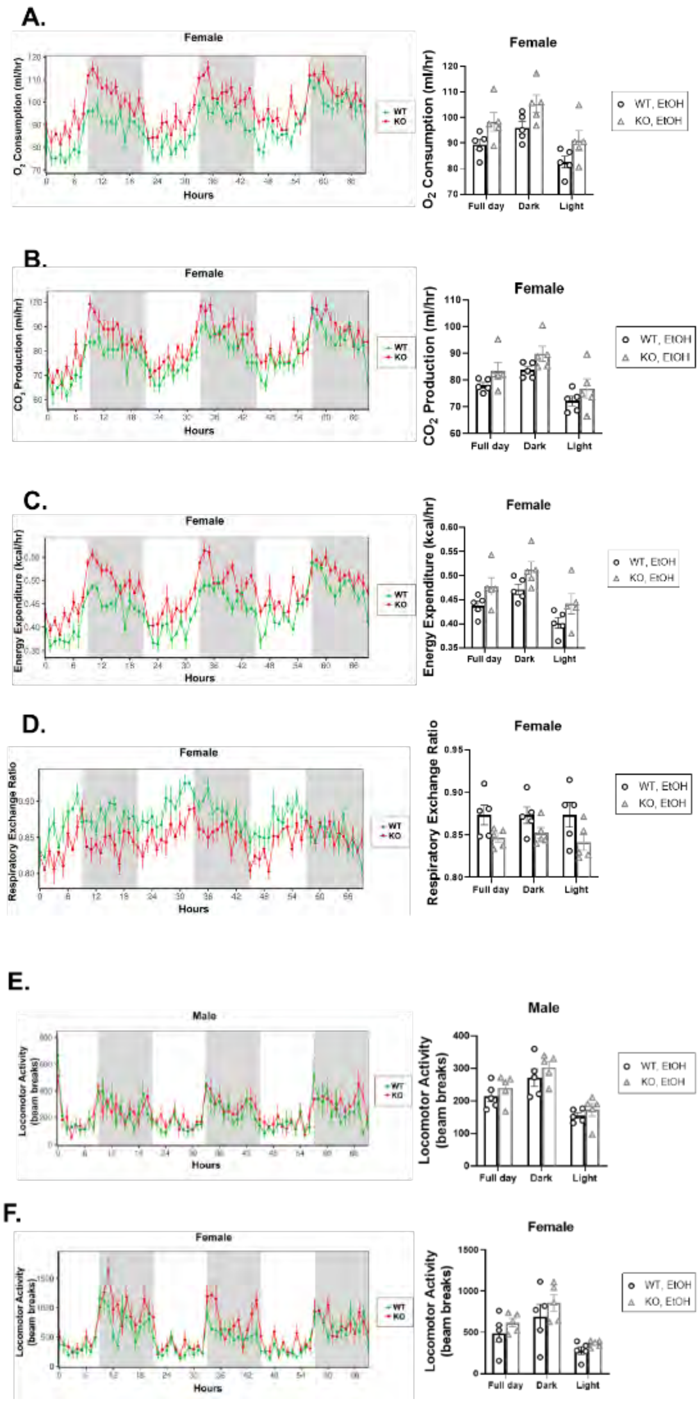

**Supplementary Figure 4**. Dihydrosphingosine (dhSph), dihydrosphingosine-1-phosphate (dhSph-1P), sphingosine (Sph), and sphingosine-1-phosphate (Sph-1P) levels in the livers of ethanol (EtOH)-fed wild-type (WT) and CerS6 knockout (KO) male and female mice. WT and CerS6 KO male and female mice were fed an EtOH diet for 6 weeks. Sphingolipid content was assessed in liver homogenates by mass spectrometry (n=5-9 per group). Data are expressed as mean ± SEM. A two-tailed unpaired *t*-test was used to compare WT and KO mice. **P* < 0.05, ***P* < 0.01, ****P* < 0.001

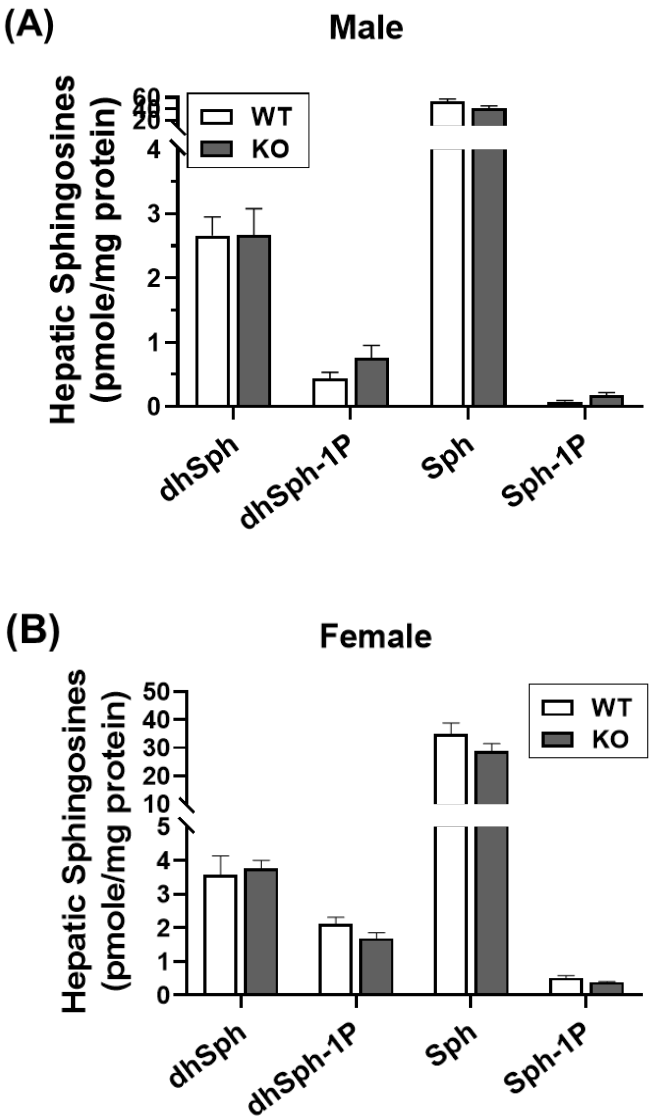

**Supplementary Table 1**. Ceramide ratios in the serum of ethanol (EtOH)-fed ceramide synthase 6 (CerS6) knockout (KO) and wild-type (WT) mice**.**

| | CerS6 KO | | | WT | | | *p*-value |
|---|---|---|---|---|---|---|---|
| **Male, Serum** | | | | | | | |
| $C_{24:0}/C_{16:0}$ | 22.69 | ± | 3.46 | 82.11 | ± | 15.24 | 0.02 |
| **Female, Serum** | | | | | | | |
| $C_{24:0}/C_{16:0}$ | 4.51 | ± | 0.31 | 66.02 | ± | 6.00 | 0.00 |

Data are reported as mean ± SEM; n=5-9/group. A two-tailed unpaired *t*-test was used to compare WT and CerS6 KO mice.
